# Mitochondrial determinants of response and resistance to venetoclax *plus* cytarabine duplet therapy in acute myeloid leukemia

**DOI:** 10.1101/2020.08.17.253856

**Authors:** Claudie Bosc, Noémie Gadaud, Aurélie Bousard, Marie Sabatier, Guillaume Cognet, Estelle Saland, Thomas Farge, Emeline Boet, Mathilde Gotanègre, Nesrine Aroua, Pierre-Luc Mouchel, Clément Larrue, Latifa Jarrou, Florian Rambow, Florence Cabon, Nathalie Nicot, François Vergez, Jérôme Tamburini, Jean-Jacques Fournié, Tony Kaoma, Jean-Christophe Marine, Christian Récher, Lucille Stuani, Carine Joffre, Jean-Emmanuel Sarry

## Abstract

The development of resistance to conventional and targeted therapy represents a major clinical barrier in treatment of acute myeloid leukemia (AML). We show that the resistance to cytarabine (AraC) and its associated mitochondrial phenotype were reversed by genetic silencing or pharmacological inhibition of BCL2 in a caspase-dependent manner. BCL2-inhibitor venetoclax (VEN) enhancement of AraC efficacy was independent of differentiation phenotype, a characteristic of response to another combination of VEN with hypomethylating agents (HMA). Furthermore, transcriptional profiles of patients with low response to VEN+AraC mirrored those of low responders to VEN+HMA in clinical trials. OxPHOS was found to be a patient stratification marker predictive of effective response to VEN+AraC but not to VEN+AZA. Importantly, whereas three cell subpopulations specifically emerged in VEN+AraC residual disease and were characterized by distinct developmental and transcriptional programs largely driven by MITF, E2F4 and p53 regulons, they each encoded proteins involved in assembly of NADH dehydrogenase complex. Notably, treatment of VEN+AraC-persisting AML cells with an ETCI inhibitor significantly increased the time-to-relapse *in vivo*. These findings provide the scientific rationale for new clinical trials of VEN+AraC combinations, especially in patients relapsing or non-responsive to chemotherapy, or after failure of frontline VEN+HMA regimen.

## Main

Despite a high rate of complete remission after conventional induction chemotherapy, overall survival is poor for acute myeloid leukemia (AML) patients, especially in elderly patients. This is due to the high frequency of relapses caused by tumor regrowth initiated by drug-resistant leukemic clones or relapse-initiating drug tolerant leukemic cells (RICs)^1^. Recent innovative therapies that are FDA-approved or under clinical development target either particular genetic features such as FLT3-ITD and IDH mutations or specific cellular processes such as BCL2 overexpression and epigenetic modifications^2–4^. Unfortunately, these new therapies do not necessarily eradicate therapeutic resistance in these patients. Indeed, recent studies have shown acquired resistance to these molecularly targeted drugs in AML^5–7^. Thus, therapy resistance remains the major therapeutic barrier in AML. Therefore, a paramount issue in the eradication of RICs is to elucidate the molecular basis of AML resistance especially *in vivo*.

While AML stem cells (LSCs) might be a chemotherapy resistant cell population of critical importance in relapse, cytarabine (AraC)-resistant persisting cells displayed high levels of reactive oxygen species (ROS), showed increased mitochondrial mass, enhanced fatty acid oxidation, and retained active polarized mitochondria, all features consistent with a high oxidative phosphorylation (OxPHOS) status^8^. Accordingly, high OxPHOS but not low OxPHOS human AML cell lines were chemoresistant *in vivo.* Targeting mitochondrial protein synthesis, electron transfer chain activities, or fatty acid oxidation (FAO) induced an energetic shift towards low OxPHOS and markedly enhanced anti-leukemic effects of AraC, suggesting a promising avenue to design new therapeutic strategies to fight AraC resistance in AML^8–11^. Mitochondrial energy metabolism and redox homeostasis also play crucial roles in response to chemotherapy in other types of hematological malignancies^12,13^ and solid tumors^14–20^.

With this perspective in mind, direct targeting of OxPHOS or inhibition of any mitochondrial function that subsequently impacts OxPHOS represents an exciting strategy to overcome drug resistance and relapses in AML. In this regard, inhibition of anti-apoptotic protein BCL-2 selectively eradicated quiescent ROS-low LSCs^21^. BCL2 suppresses apoptosis in a variety of cell types and regulates cell death by controlling the mitochondrial membrane permeability through a feedback loop system with caspases. This inhibits caspase activity by preventing the release of cytochrome c from mitochondria. Preclinical data demonstrated the anti-leukemic efficacy of the BCL2 inhibitor venetoclax (VEN) in AML and its synergy when combined with hypomethylating or chemotherapeutic agents^22–24^. Recent clinical trials have confirmed the clinical benefit of these VEN-based therapies in newly diagnosed AML, leading to the recent FDA approval of VEN in combination with hypomethylating agents (HMA; azacitidine, AZA or decitabine) or low-dose AraC for older adults with newly diagnosed AML ^25^. Several mechanisms of acquired resistance to VEN alone or VEN+AZA in AML have been identified and highlighted the crosstalk between mitochondrial metabolism and survival pathways in AML. However, it remains to be clarified how BCL2 affects OxPHOS (potentially by preventing the loss of cytochrome c from the intermembrane space through mitochondrial outer membrane permeabilization (MOMP)) and whether it is involved in the resistance to various VEN-based therapies. Furthermore, mechanisms of action/resistance specific for the VEN+AraC combination have also yet to be elucidated.

To specifically address these questions and identify the mechanism of resistance responsible for the relapse in patients treated with this combination, we have developed treatment strategies targeting the BCL2 dependency of AraC resistant cells *in vivo* using our NOD/LtSz-scid IL2Rγ_c_^null^ (NSG) immunodeficient mice-based patient-derived xenograft (PDX) models. Here, we first demonstrated that VEN enhances the efficacy of AraC *in vitro* and *in vivo* by reversing hyperactivation of OxPHOS observed upon AraC treatment in a caspase-dependent and differentiation status-independent manner as opposed to the VEN+AZA combination. Unexpectedly, we discovered that mitochondrial electron transport chain complex I (ETCI) organization and activity are involved in residual disease post-VEN+AraC *in vivo*. Treatment with selective potent ETCI inhibitor IACS-010759 for 9 weeks in consolidation phase following VEN+AraC significant enhanced time-to-relapse compared to the placebo arm, suggesting promising new combinatory therapeutic strategies in AML.

## Results

### Mitochondrial permeability transition pore and mitochondrial BCL2 are involved in OxPHOS-dependent resistance of AML cells to cytarabine

Previous studies showed that residual AML cells with high OxPHOS activity prevent mitochondrial priming to cell death and apoptosis^8,26–28^. Accordingly, they have an increased mitochondrial membrane potential (MMP) without any changes in mitochondrial mass post-AraC ^8^, indicating that retained active polarized mitochondria were maintained at relapse (Supplementary Fig. S1A-S1B). These data confirmed the role of the MMP in AML cells not only at the nadir but also at relapse. Indeed, AraC-induced mitochondrial apoptotic cell death is due to a sustained mitochondrial depolarization caused by increased ROS-induced MOMP and mitochondrial priming of apoptosis in AML cells^8,29–31^. In this context, a mitochondrial inner transmembrane protein pore (mitochondrial permeability transition pore, mPTP), is normally closed but when opened allows passage of small (less than 1.5□kDa in mass) molecules, it may lead to release of apoptotic factors if and when the outer mitochondrial membrane is breached^32^. Furthermore, mPTP and MOMP also play an important role in mitochondrial metabolite and Ca^2+^ release/accumulation through the inner mitochondrial membrane^33–37^, thus affecting both TCA cycle and OxPHOS activity and capacity ^38–40^. Therefore, we proposed that this may in part due to the bidirectional interplay between MOMP and the mPTP opening in AML.

To confirm this link, we first analyzed mPTP opening/closing and mitochondrial calcium content using the CoCl_2_-calcein fluorescence-quenching assay by flow cytometry^41,42^ (Fig. 1A) and the rhod-2 fluorescent probe, respectively in viable AML cells after PBS- and AraC- treatment. The mitochondrial retained calcein fluorescence is commonly used as a proxy to denote opening/closing of the mPTP, while Rhod-2 is as a selective mitochondrial Ca^2+^ fluorescent sensor. In presence of the calcein fluorescence quencher CoCl2, the fluorescence of cytosolic calcein is lost, while mitochondrial calcein fluorescence is retained. We assessed mPTP opening/closing in nine patient-derived xenograft (PDX) and one cell line-derived xenograft (CLDX) in an immunodeficient (NOD/LtSz-scid IL2Rγ_c_^null^, NSG) mouse model^43^ (Supplementary Table S1). We found that calcein retention was increased in the presence of CoCl2 in residual AML cells post-AraC *in vivo*, suggesting an inhibition of mPTP opening (Fig. 1A). Moreover, the extent of calcein retention, *i.e.* inhibition of mPTP opening by AraC compared to PBS, correlated with the sensitivity to AraC *in vivo* (Fig. 1B). PDXs with the greatest sensitivity to inhibition of the mPTP by AraC (*e.g.* the highest mitochondrial priming to apoptosis) are the strongest responders to AraC *in vivo* (Fig. 1B-1C). We further confirmed this observation *in vitro* in residual AML cells post-AraC (Fig. 1D) and, in line with this, we found that mitochondrial Ca^2+^ is also significantly increased in viable AML cells post-AraC (Fig. 1E). These observations in conjunction with our previous published work^8^ linked MMP, mPTP activity and mitochondrial matrix Ca2+ to OxPHOS activity and survival of AraC residual cells *in vitro* and *in vivo*.

**Figure 1.**
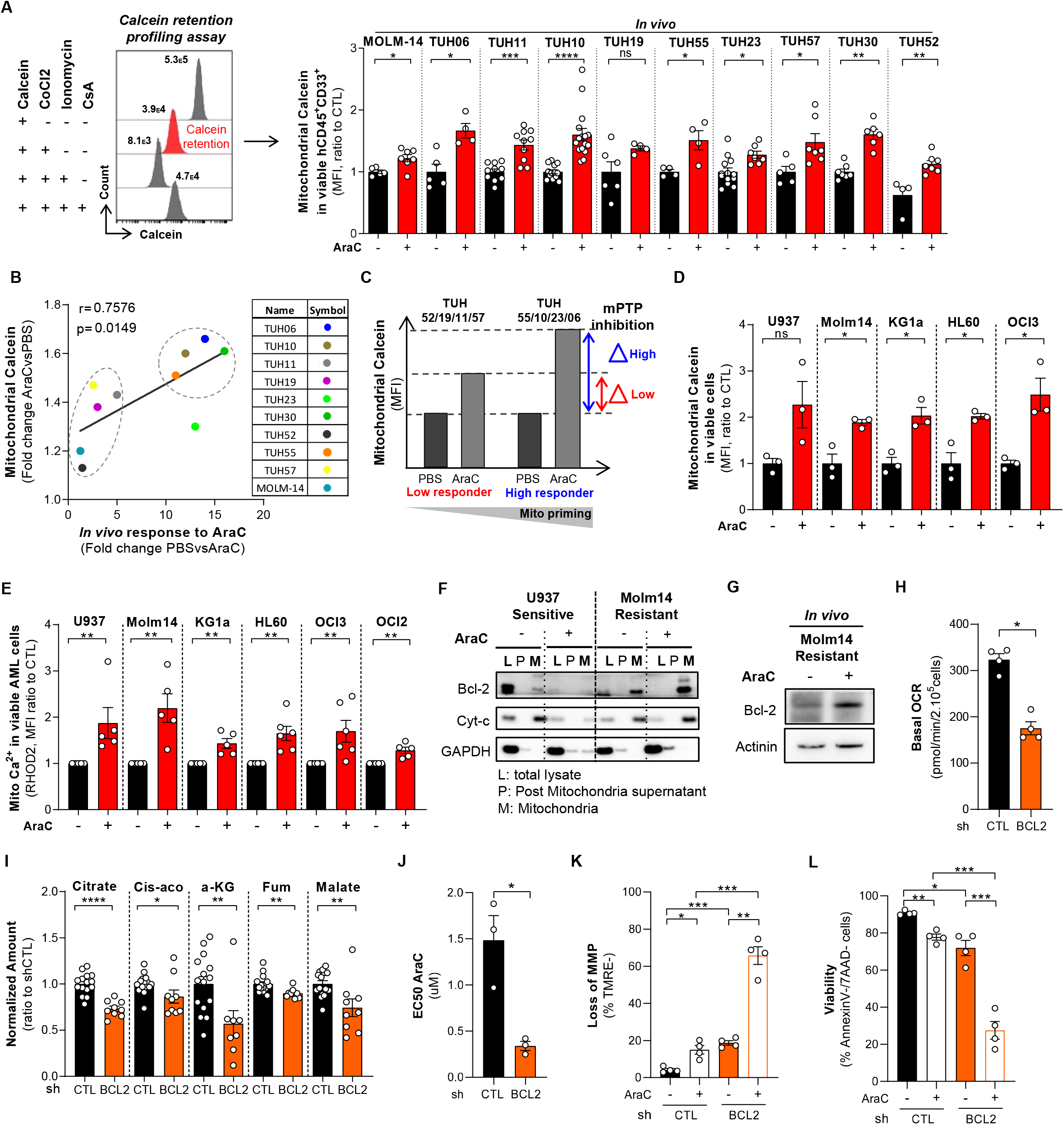
Identification of mitochondrial permeability transition pore as a key actor of early resistance to cytarabine in AML. **(A)** The opening of mitochondrial permeability transition pore (mPTP) in AML cells was measured with the calcein/cobalt quenching assay. The calcein retention profiling assay measures the calcein retained into mitochondria following the addition of CoCl2. The calcium ionophore ionomycin (2.5 μM) was used as control of the mPTP opening while cyclosporine A (10 μM) was used as control of closed mPTP. The fluorescence from cytosolic calcein was quenched by the addition of CoCl2, while the fluorescence from the mitochondrial calcein was maintained. The MFI of the mitochondrial retrained calcein fluorescence from primary AML blasts following *in vivo* AraC treatment in 9 different PDXs (60 mg/kg) and 1 CLDX (MOLM14, 30 mg/kg) is shown. Each dot is a mouse from the control or AraC group. Values are mean ± SEM. **(B)** The fold change in mitochondrial retrained calcein of viable humain AML blasts after *in vivo* AraC treated group vs the control group correlates with the *in vivo* response to AraC. Each dot is the mean of a single *in vivo* PDX or CLDX experiment. **(C)** The fold change in retained calcein after *versus* before AraC treatment was called “mPTP inhibition”. Indeed, patient good responders to AraC in vivo (TUH55, TUH10, TUH23, TUH06) have a higher mPTP inhibition, meaning that they are more “primed” to apoptotic cell death than low responders to AraC (TUH52, TUH19, TUH11, TUH57). **(D)** Mitochondrial retained calcein fluorescence in viable AML cells before and after AraC treatment (2μM) *in vitro* in 6 different AML cell lines. Each value is an independent experiment, N=3. Values are mean ± SEM. **(E)** Mitochondrial calcium content after *in vitro* AraC treatment (2μM) was obtained using the rhodamin2 probe *in vitro* in 6 different AML cell lines. Each value is an independent experiment, N=5 or 6. Values are mean ± SEM. **(F)** Mitochondrial preparation after *in vitro* AraC treatment (2μM) from AraC-sensitive cells (U937) or from AraC-resistant cells (MOLM14). Bcl-2 is present at mitochondria in basal condition only in resistant cells and remain at mitochondria after AraC treatment. **(G)** Bcl-2 expression in MOLM14 AML cells after *in vivo* AraC treatment (30mpk) in NSG mice. **(H)** MOLM14 transduced with CTL or BCL2 shRNAs were examined for their basal oxygen consumption rates. Each value is an independent experiment, data are mean ± SEM, (N=4). **(I)** Citrate, cis-aconitate (Cis-aco), α-ketoglutarate (α-KG), fumarate (Fum) and malate amounts measured by IC/MS in MOLM14 cells transduced with CTL or BCL2 shRNAs. Data are mean ± SEM. Each dot is an experimental replicate of three independent experiments. **(J)** EC50 for AraC of MOLM14 transduced with CTL or BCL2 shRNAs analyzed after 24h of treatment using annexinV/7AAD flow cytometry staining. Data are mean ± SEM, (N=3). **(K,L)** Loss of mitochondrial membrane potential (K) assessed by flow cytometry using fluorescent TMRE probe staining and percent of viable cells (L) assed by annexinV/7AAD flow cytometry staining following 24H of AraC treatment (2μM) in MOLM14 transduced with CTL or BCL2 shRNAs. Data are mean ± SEM, (N=4).

Interestingly, viable AML blasts with increased MMP without a change in mitochondrial mass (Supplementary Fig. S1A-S1B) displayed an increased BCL2 expression at relapse post-AraC treatment (Supplementary Fig. S1C). Furthermore, AML blasts have a strong dependency on BCL2 and exhibit higher BCL2 gene and protein expression levels compared to normal cells (Supplementary Fig. S1D-S1E), suggesting key biological roles for this protein in AML by modulating both mitochondrial metabolic and apoptotic processes^22,29^. To determine whether BCL2 affects the mPTP, mitochondrial Ca^2+^ homeostasis and mitochondrial priming to cell death in AML, we investigated the expression and the role of BCL2 in our AraC resistance model. First, we observed that AraC-resistant high OxPHOS MOLM14 AML cells overexpressed total and mitochondrial-bound BCL2 *in vitro* but not in AraC-sensitive U937 cells (Fig. 1F). We also quantified the expression of BCL2 in human viable AML cells purified from mice xenografted and treated with AraC *in vivo*. Similarly, BCL2 expression was increased in AraC-residual MOLM14 cells from bone marrow (Fig. 1G). Since BCL2 overexpression is associated with high OxPHOS state and AraC resistance in AML, we analyzed the impact of the silencing of BCL2 in AraC-resistant MOLM14 cells (Supplementary Fig. S1F). We found that BCL2 silencing in AraC-resistant MOLM14 cells decreased the basal rate of mitochondrial oxygen consumption (Fig. 1H) and the level of TCA cycle intermediates (Fig. 1I). This significantly sensitized cells to AraC (Fig. 1J) through the enhancement of loss of MMP (Fig. 1K) and a marked reduction of cell viability *in vitro* (Fig. 1L). We consistently observed that AML patients with the highest BCL2 expression from our TUH and the BeatAML^44^ cohorts were the most sensitive to BCL2 inhibitor venetoclax (VEN) (Supplementary Fig. S1G-H). Furthermore, they displayed an enrichment in gene signatures related to leukemic stem cells, cysteine metabolism or mitochondria (Supplementary Fig. S1H-S1J), all known markers of response to either VEN or VEN+AZA^25,45–47^. Finally, transcriptomes of VEN-resistant AML patients are enriched in gene sets related to hypoxia and glycolysis.

In summary, BCL2 is critical for optimal OxPHOS activity of AraC resistant AML cells through the inhibition of the mPTP (Supplementary Fig. S1K). These data confirmed and extended previous studies, showing that BCL2 modulates both mitochondrial OxPHOS and mitochondrial priming of AML cells to genotoxic-induced apoptosis^8,21,22,29,30^. Together, we further demonstrate that AraC promotes BCL2 overexpression *in vitro* and *in vivo*, its localization at mitochondria and that this process is associated with elevated mitochondrial calcium, mitochondrial respiration, TCA cycle metabolites and mPTP closure state.

### BCL2 inhibitor venetoclax improves anti-leukemic activity of cytarabine in a differentiation dependent manner

Consistent with the above-mentioned results, we speculated that the highly specific BCL2 inhibitor venetoclax (VEN or ABT-199) might enhance anti-AML effects of AraC. To end this, we first assessed the combinatory effect of AraC and VEN *in vitro* with three AML cell lines expressing variable BCL2 levels and mutational status (Supplementary Fig. S1E; Supplementary Table S2). We observed a synergistic effect between AraC and VEN in these cell lines with the strongest, intermediate and lowest synergy^48^ in MOLM-14, U937 and OCI-AML3 cells, respectively (Supplementary Fig. S2A-S2F) through a reduction of cell viability, the dissipation of MMP and the induction of apoptosis (Supplementary Fig. S2G-S2H).

Next and importantly, we developed a duplet combinatory therapeutic strategy targeting the BCL2 dependency of AraC resistant AML cells *in vivo* using our PDX models to mimick the therapeutic regimen used in clinical trials of VEN *plus* AraC for AML treatment (Fig. 2A). We injected primary AML patient specimens into the tail vein of mice and when the disease was established, we treated for one week with AraC (30mg/kg/d) with or without VEN (100mg/kg/d). Induction of apoptosis (loss of MMP) and reduction in total cell burden in spleen (SP) and bone marrow (BM) were quantified in each cohort at the day after the last dose. As expected, VEN improved anti-AML activity of AraC through an enhanced dissipation of MMP *in vivo* in six different PDX models (Fig. 2B; Supplementary Fig. S3A). While 3 of the 6 responses were significant with VEN+AraC compared to AraC, all of these PDX had a marked increase in fold reduction (ranged from 19 to 410-fold) post-VEN+AraC compared to post-AraC. Of note, this was particularly true in lowest responders to AraC (fold reduction post-AraC lower than 10). Importantly, the effects on body weight, normal hematopoietic cells and hematological parameters upon combination therapy were primarily driven by AraC toxicity (Supplementary Fig. S3B-S3D).

**Figure 2.**
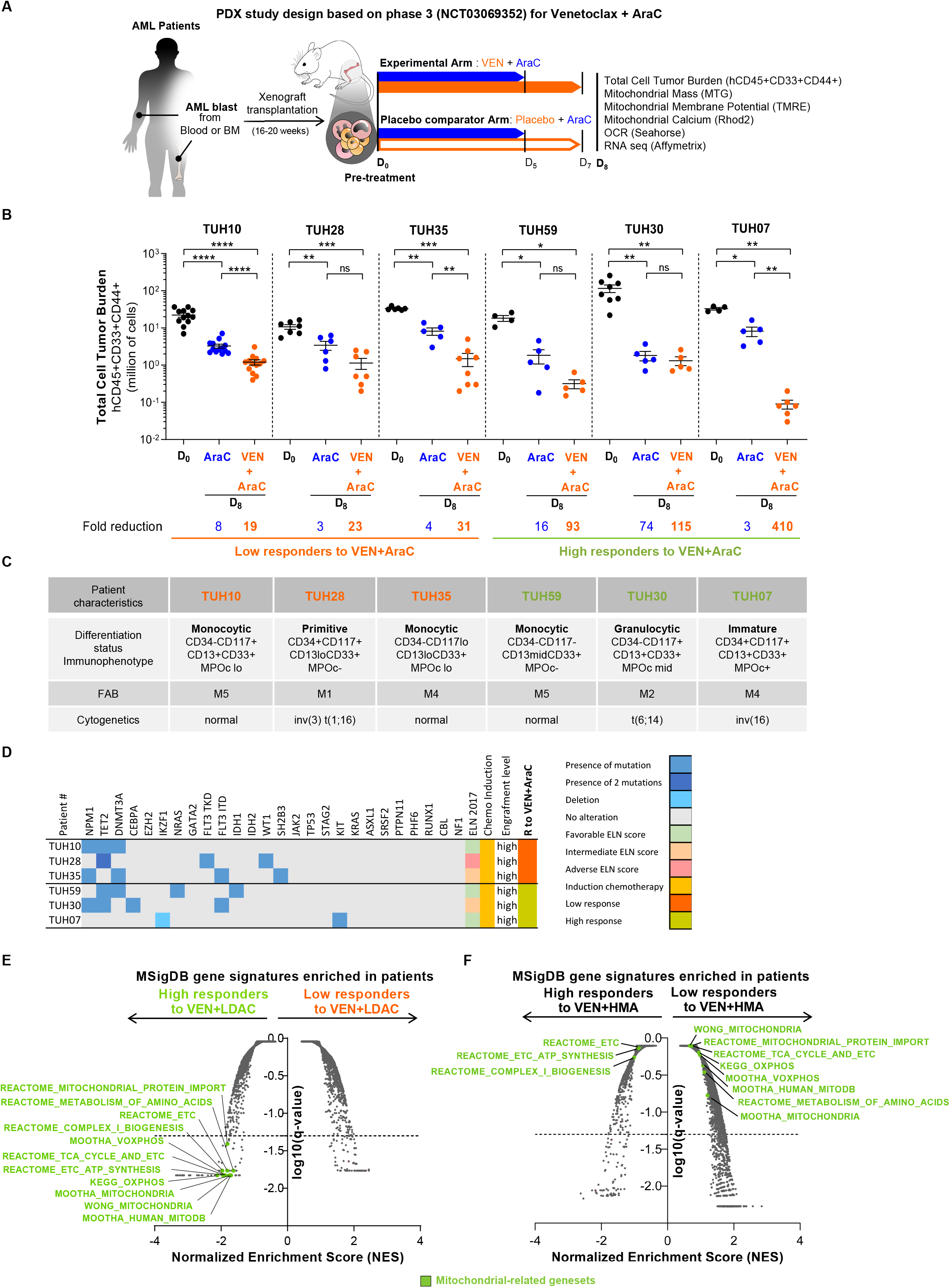
Venetoclax improves anti-tumor activity of cytarabine with mechanisms-of-action/resistance different from venetoclax plus HMA in AML PDX models and patients. **(A)** Schematic diagram of the therapeutic regimen schedule based on the phase 3 venetoclax (VEN) plus cytarabine (AraC) (NCT03069352) in NSG mice xenografted with patient AML samples. The experimental arm received 30 mg/kg/day of AraC given daily *via* intraperitoneal injection for 5 days and 100 mg/kg/day of venetoclax given daily *via* oral gavage for 7 days. **(B)** Cumulative total cell tumor burden (TCTB) of human viable CD45^+^CD33^+^CD44^+^ AML cells assessed in bone marrow and spleen at day 0 (D_0_) and at day 8 (D_8_) for the experimental arm or the placebo comparator arm by flow cytometry in 6 different PDXs. Each dot is a mouse. Values are mean ± SEM. **(C)** Summary of patient characteristics (immunophenotype, FAB classification and cytogenetics), ordered based on the in vivo response to the duplet therapy venetoclax plus AraC. **(D)** Concomitance of frequently mutated genes in AML and genetic risk classification “favorable”, “intermediate” and “adverse” based on the European Leukemia Net (ELN). Each line represents 1 of 6 patient samples used for PDX model, ordered based on the *in vivo* response to the duplet therapy venetoclax plus AraC. **(E,F)** Gene set enrichment analysis (GSEA) of gene signatures from the Broad Institute Molecular Signature Data Base (MSigDB) at diagnosis in patients receiving the duplet therapy venetoclax plus low dose acracytine (LDAC) (E) or hypomethylating agent (HMA) (azacitidine or decitabine) (F). Dotted line at −1.3 (log10(0.05)) indicates the threshold limit of the q-value below which the gene signatures are significantly enriched.

Finally, to identify putative markers predictive of the response to this duplet therapy, we stratified our PDX in two subgroups based on their response (fold reduction; cut-off: 60) to VEN+AraC (Fig. 2B-2C). Interestingly and as opposed to previously reported treatment with VEN+AZA^49^, none of the examined features was associated with myeloid differentiation status (such as CD34/CD117/CD13/CD33/MPO-based immunophenotype, NPM1 status and FAB classification system) or segregated AML patients for their own *in vivo* response (Fig. 2C). Mutational status, cytogenetics and ELN2017 score also did not discriminate low and high responders to VEN+AraC: (Fig. 2D; Supplementary Table S3-S4). More importantly, we analyzed gene expression from 31 patients receiving VEN combined with hypomethylating agents (HMA, AZA) or low-dose cytarabine (LDAC, AraC) and stratified as high (CR, complete remission and durable remission) or low (RD, refractory disease) responders to these tow VEN-based combinations ^25^. This revealed that gene signatures related to mitochondrial metabolism and ETC biogenesis are enriched in high responders to VEN+LDAC (Fig. 2E and supplementary Fig. S4A). Unexpectedly, this is different from high and low responders to VEN+HMA compared to high and low responders to VEN+LDAC (Fig. 2F and supplementary Fig. S4B). These data indicate that predictive markers of VEN+AraC are different from VEN+AZA, suggesting that the molecular mechanisms of both action and resistance to these two VEN combinations are likely different. Of note, AML patients with a high OxPHOS phenotype/signature would be selectively high responders to VEN+AraC. Indeed, there are independent of differentiation status and more depend on specific metabolic status leading to mitochondrial organization and OxPHOS hyperactivation in AML.

### AraC-induced resistance and bioenergetic capacity are rescued by the duplet therapy VEN+AraC in a calcium- and caspase- dependent manner

shBCL2 (Fig. 1) or BCL2 inhibitors (navitoclax in AML^21^; VEN: in breast cancer^50^, in lymphoma^51^ and in AML^52^) reduce mitochondrial respiration. Similarly, VEN+AZA targets LSCs by inhibiting mitochondrial OxPHOS^47,53^. Consistently, we next sought to test whether VEN+AraC also affects mitochondrial respiration and metabolism in AML cells. While VEN marginally tended to decrease the basal rate of mitochondrial oxygen consumption (OCR) compared to control, VEN markedly inhibited the increase in OCR induced by AraC *in vitro* and *in vivo* (Fig. 3A-3B; Supplementary Fig. S5A). This is not due to a reduction of mitochondrial mass and ETC subunit protein expression *in vitro* and *in vivo* (Supplementary Fig S5B-S5D). These results suggest that this drug combination instead inhibited the activity of ETC complexes or/and availability of substrates or cofactors required for OxPHOS/ETC activity rather than their expression levels. Accordingly, VEN abrogated the increased ETCI activity and several key TCA cycle intermediates observed upon AraC treatment (Fig. 3C-3D). VEN also abolished the enhancement in levels of glycolytic and pentose phosphate pathway intermediates in response to AraC (Supplementary Fig. S5E-S5G). Furthermore, VEN significantly blocked the induction of lactate production by AraC (Fig. 3E). Altogether these results suggest the combination of VEN+AraC affects glycolytic flux to support cellular bioenergetic metabolism in AML. To assess the contribution of glycolysis to ATP production in AML cells, we calculated the glycolytic ATP production rate (J_ATP_GLY) from ECAR measurements using the methodology developed by Mookerjee and Brand^54^. Accordingly, VEN+AraC treated cells displayed a significantly lower J_ATP_gly compared to AraC treated cells (Fig. 3F). We next examined the bioenergetic capacity of AML cells by measuring maximal OxPHOS and glycolysis ATP production rates. The maximal JATPox was determined using the mitochondrial uncoupling agent FCCP and the maximal JATPgly was determined using the ionophore monensin, which stimulates plasma membrane Na+/K+ pumps to drive increased ATP demand under conditions of mitochondrial inhibition^54^. VEN lowered both basal and AraC-stimulated bioenergetic capacity in AML cells (Fig. 3G). Hence, we demonstrate that targeting BCL2 with VEN abrogates enhanced global cellular bioenergetic capacity of AraC resistant AML cells. This prevents the induction of a compensatory Pasteur effect in response to a decrease in OxPHOS reduction elicited by VEN in the presence of AraC. In fact, both ETC1 and OxPHOS activities were maintained at the same level as those in control cells, suggesting that residual cells did not undergo an energetic crisis and were able to support their energetic/ATP demand to survive.

**Figure 3.**
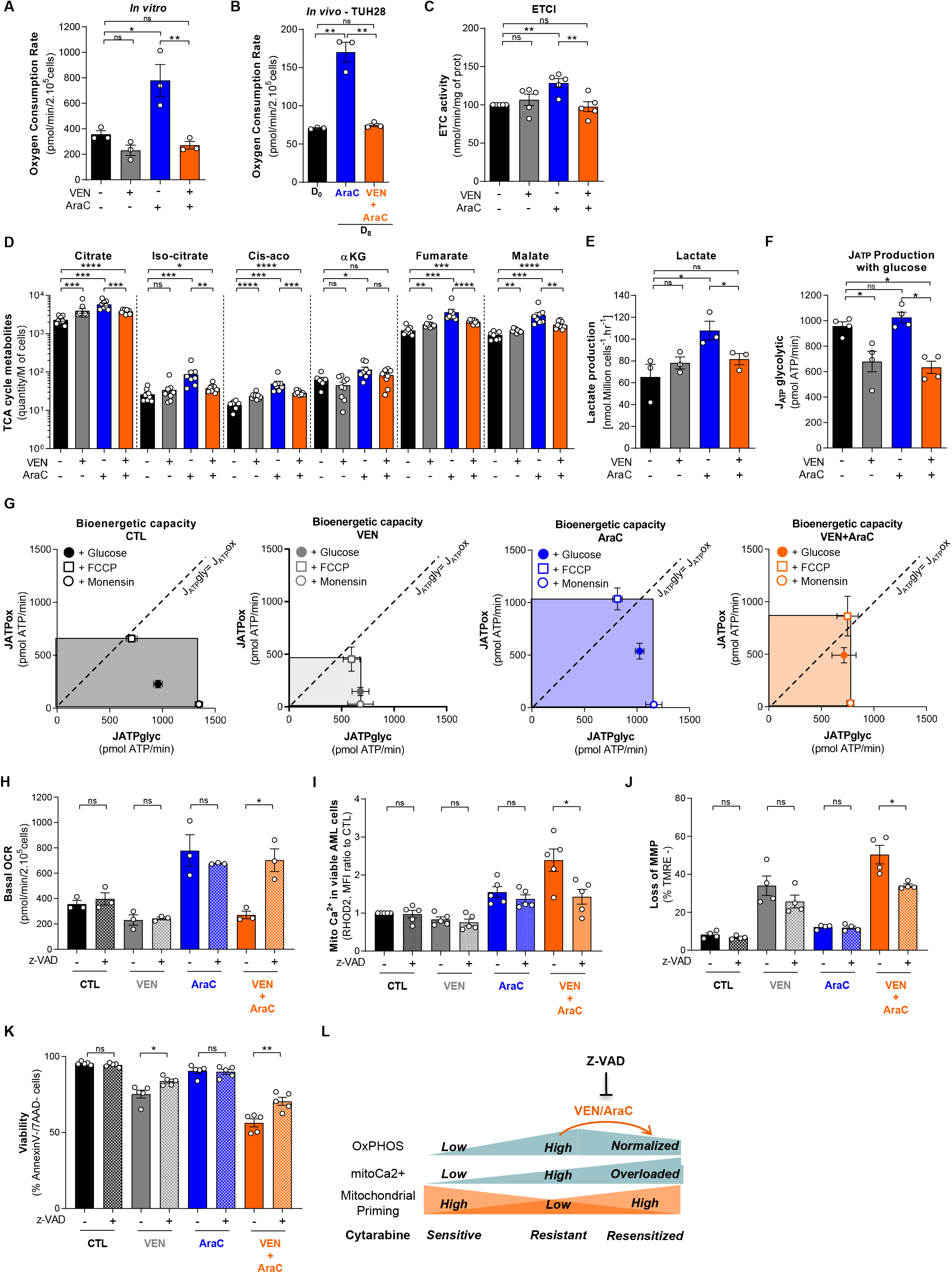
AraC-induced resistance and bioenergetic capacity are rescued by the duplet therapy VEN+AraC in a calcium- and caspase-dependent manner. **(A)** Histogram shows the basal oxygen consumption rate measured by Seahorse XF24 Extracellular Analyzer of *in vitro* MOLM14 cells treated or not with venetoclax (VEN, 0.5 μM) and cytarabine (AraC, 0.5 μM) for 24H (N=3). Values are mean ± SEM. **(B)** Basal oxygen consumption rate of PDX TUH28 assessed by Seahorse XF24 Extracellular Analyzer in FACS-sorted human AML cells harvested from mice at day 8, following the end of the therapy. **(C)** Mitochondrial ETC complex I activity in MOLM14 cells following 24H treatment with venetoclax (VEN, 0.5 μM) and cytarabine (AraC, 0.5 μM) (N=5). Values are mean ± SEM. **(D)** Citrate, iso-citrate, cis-aconitate (Cis-aco), alpha-ketoglutarate (α-KG), fumarate and malate amount measured by IC/MS in MOLM14 cells following 24H treatment with venetoclax (VEN, 0.5 μM) and cytarabine (AraC, 0.5 μM). Amounts are in pmol/M of cells, except α-KG and fumarate which are in ratio (12C/13C)*1000. Data are mean ± SEM. Each dot is an experimental replicate of three independent experiments. **(E)** Lactate amounts measured by NMR in extracellular medium of MOLM14 cells following 24H treatment with venetoclax (VEN, 0.5 μM) and cytarabine (AraC, 0.5 μM). Data are mean ± SEM (N=3). **(F)** Quantification of glycolytic ATP production (JATPglycolytic) following 10mM glucose addition for MOLM14 cells treated for 24H with venetoclax (VEN, 0.5 μM) and cytarabine (AraC, 0.5 μM) after the addition of glucose (10 mM), using a Seahorse XF24 Extracellular Analyzer (N = 4). Values are mean ± SEM. **(G)** Bioenergetic capacity plots of MOLM14 cells following treatment with venetoclax (VEN, 0.5 μM) and cytarabine (AraC, 0.5 μM) for 24H. Points in the bioenergetic space represent glycolytic ATP production (JATPglyc) and oxidative ATP production (JATPox) for 2×10^5^ cells following addition of glucose (full circle), FCCP (empty square), and monensin (empty circle) (N = 4). **(H)** Basal oxygen consumption rate in MOLM14 cells following treatment with venetoclax (VEN, 0.5 μM) and cytarabine (AraC, 0.5 μM) in presence or not of z-VAD (40 μM) for 24H. Values are mean ± SEM (N=3). **(I)** Mitochondrial calcium content in MOLM14 cells following treatment with venetoclax (VEN, 0.5 μM) and cytarabine (AraC, 0.5 μM) in presence or not of z-VAD (40 μM) for 24H, assessed by flow cytometry using the rhodamin2 probe. Values are mean ± SEM (N=5). **(J)** Percent of loss of mitochondrial membrane potential in MOLM14 cells following treatment with venetoclax (VEN, 0.5 μM) and cytarabine (AraC, 0.5 μM) in presence or not of z-VAD (40 μM) for 24H, assessed by flow cytometry using the TMRE probe. Values are mean ± SEM (N=4). **(K)** Percent of viable MOLM14 cells following treatment with venetoclax (VEN, 0.5 μM) and cytarabine (AraC, 0.5 μM) in presence or not of z-VAD (40 μM) for 24H assessed by annexinV/7AAD flow cytometry staining. Values are mean ± SEM (N=5). **(L)** Schematic diagram of the mechanisms-of-action of VEN+AraC combination that reversed the induction of mitochondrial phenotype by AraC leading to mitochondrial priming of the AraC residual/resistant cells to cell death. These mechanisms are abrogated by a pan-caspase inhibitor Z-VAD-fmk.

To investigate the mechanism of cell death induced by this duplet therapy VEN+AraC, we treated AML cells exposed to VEN+AraC or single agents with the irreversible pan-caspase inhibitor Z-VAD-FMK that prevents apoptosis by blocking initiator caspases (such as caspase-2, −8, −9, and −10) and executioner/effector caspases (caspase-3, −6, and −7)^55^. We showed that Z-VAD-FMK rescued not only OCR, but also mitochondrial calcium, loss of MMP and cell viability selectively upon VEN+AraC treatment (but not upon single cell treatments; Fig. 3H-3K). This indicates that Z-VAD-FMK might suppress BCL2-dependent Ca2+ overload and cell survival, and that caspases may act as inhibitors of mitochondrial priming to apoptosis in this BCL2 model of AML chemoresistance (Fig. 3L).

### Minimal residual disease from duplet therapy VEN+AraC is associated with cell clusters exhibiting distinct developmental and metabolic programs

Next, we characterized mitochondrial function in residual disease post-VEN+AraC. We did not observe any significant difference in mitochondrial membrane potential in viable AML cells after the duplet combination compared to single agents *in vitro* and *in vivo* (Supplementary Fig. S6A-S6B). This may indicate that residual cells are able to maintain mitochondrial activity in response to stress induced by the duplet therapy VEN+AraC.

To further decipher specific mechanisms of tolerance and subsequent resistance to the combination of VEN+AraC compared to single agents, we analyzed transcriptomes of more than 15K single cells (scRNAseq; 10x genomics, Supplementary Figure S7A) from purified viable AML blasts of PDX model of a high responder patient TUH07, before treatment, after VEN or AraC treatments alone, or after combination of both (VEN+AraC) (Fig. 4A). We first confirmed that duplet combination was more effective in reducing total disease burden than AraC alone or VEN alone (Supplementary Figure S7B). Consistent with our previous work^8^, these experiments confirmed the emergence of cells displaying a gene signature enriched in fatty acid translocase CD36 under AraC treatment at the single cell level (Supplementary Fig. S7C). While performing Seurat and unsupervised hierarchical clustering analysis on all cell transcriptomes, 15 distinct transcriptional states were detected (Fig. 4B; Supplementary Fig. S7D). The proportion and presence of cells in those states varied between conditions, especially after duplet therapy (Fig. 4C-4D). Notably, the proportion of cells belonging to clusters #0, #1, and #11 was increased in both AraC and VEN-residual disease compared to vehicle-treated AML cells. Those clusters were enriched in gene sets related to key biological processes including immune response, cell differentiation, migration, DNA repair and epigenetics (Supplementary Fig. S7E). Furthermore, the cell proportion of clusters #10, #7 and #8 increased in AML cells treated with AraC alone and VEN alone, respectively (Fig. 4C-4E). Cell clusters specific to VEN resistance (#7, #8) were notably enriched in gene sets related to calcium ion homeostasis (Supplementary Fig. S7E), confirming the relationship between BCL-2 and Ca^2+^ regulation.

**Figure 4.**
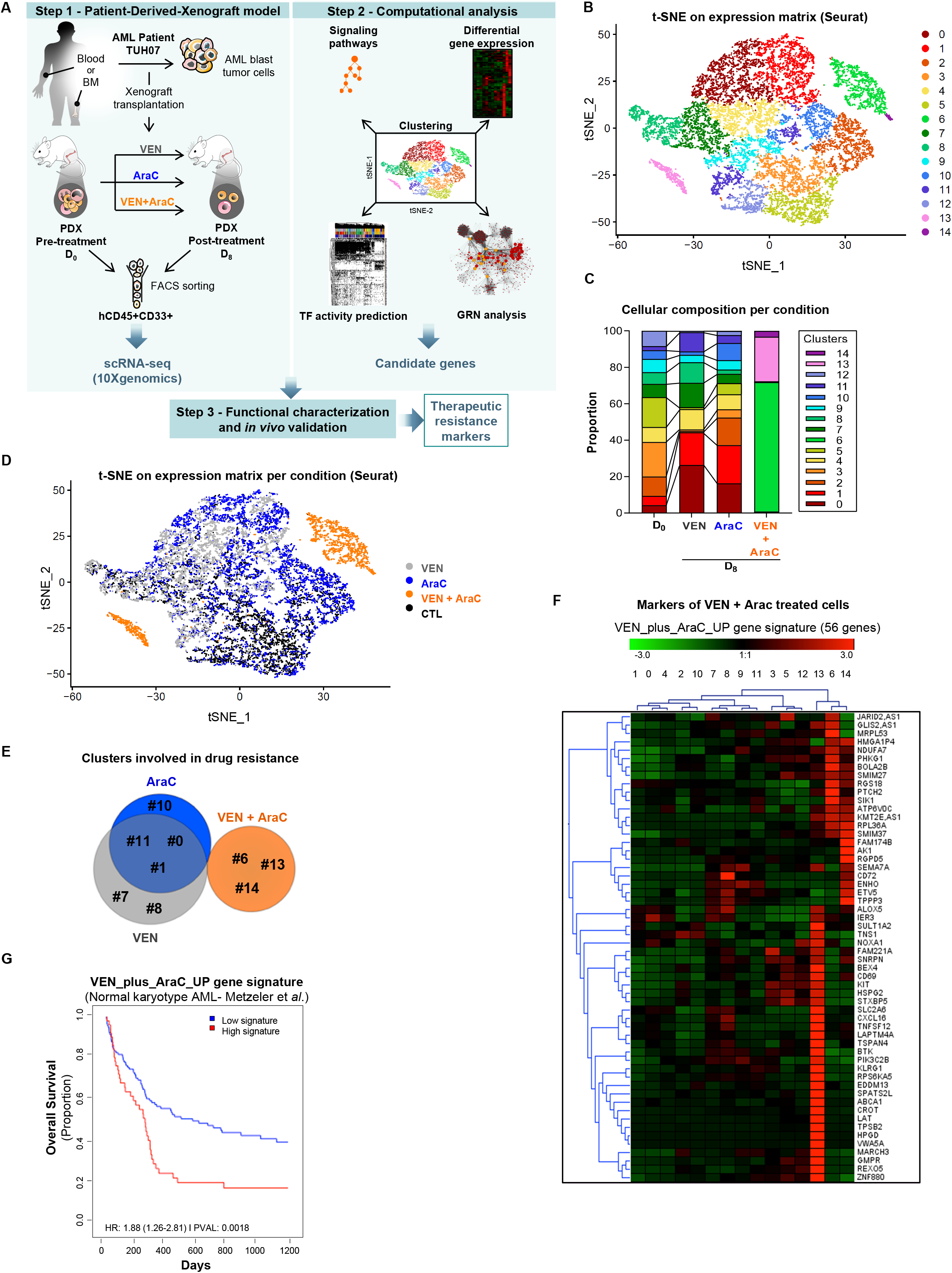
Single-cell RNA-seq analysis uncovers cell states specific of the duplet-therapy resistance. **(A)** Schematic representation of the experimental strategy. *In vivo* AML residual cells were harvested from PDX TUH07 before and after three different treatments: monotherapy venetoclax (VEN), monotherapy cytarabine (AraC) and the duplet therapy (VEN+AraC). Human myeloid cells (CD45^+^CD33^+^) were sorted using FACS, and single-cell transcriptomes were generated using 10X technology. Single cells were clustered using Seurat. Differential gene expression analysis, pathway enrichment analysis, transcription factor (TF) activity prediction and gene regulatory network (GRN) analysis were used to identify candidate genes involved in therapeutic resistance. Finally, a functional characterization of the candidate genes leads to therapeutic resistance markers discovery. **(B)** t-SNE plot of the transcriptomes of 15,604 sequenced single cells in the four conditions combined (Day0; Day8 following VEN; AraC; VEN+AraC) using Seurat. Colors indicate k-means clusters (k□=□15). **(C)** Proportion of cluster repartition per condition (Day0; Day8 following VEN; AraC; VEN+AraC). **(D)** t-SNE plot colored by treatment condition. **(E)** Schematic representation of clusters involved in drug resistance under AraC (blue), VEN (grey), VEN+AraC (orange) treatments. The area represented in each color is in proportion to the number of single cells in the corresponding color of the Venn diagram. **(F)** Heatmap showing the gene set called VEN_plus_AraC_UP gene signature, overexpressed in at least one of the three clusters appearing following the duplet therapy VEN+AraC compared to other clusters preexisting before treatment. **(G)** The VEN_plus_AraC_UP gene signature expression predicts the survival rate response from Metzeler dataset in normal karyotype AML.

Unexpectedly, VEN+AraC residual AML cells constituted a group of three cell clusters, #6, #13 and #14, that were not present in residual AML blasts following treatment with either AraC alone or VEN alone (Fig. 4D-4E). This suggests the emergence of multiple and duplet-specific transcriptional states after VEN+AraC. Furthermore, no cells from pre-treatment samples were present in these three clusters, indicating that this emergence was not a selective but an adaptive response specific to the duplet treatment *in vivo.* Further gene set enrichment analysis of the GO terms identified the up-regulation of OxPHOS activity, ETC assembly, and mitochondrial organization in clusters #6 and #14 (Supplementary Fig. S7E). OxPHOS signature was also observed as enriched in cluster #13, even at a lesser extend (Supplementary Fig. S8A). GO analysis revealed that cluster #13 was enriched in genes involved in lipid/FA metabolism, angiogenesis and immune response (Supplementary Fig. S7E; S8B). Interestingly, based on Van Galen hierarchies^56^, these three emerging cell subpopulations were enriched in gene signatures previously reported as defining HSPC-like, promonocyte-like or the cDC-like myeloid developmental stages, respectively (Supplementary Fig. S8C). Unsupervised clustering of all genes upregulated in these three cell clusters compared to all other clusters identified a specific VEN+AraC resistance gene signature (VEN_plus_AraC_UP-gene signature) encompassing 56 genes (Fig. 4F, Supplementary Fig. S8D). Datamining analyses of this specific gene signature using Genomatix showed that it is enriched in genes for lipid metabolic processes (Supplementary Fig. S8E). When the independent Metzeler cohort^57^ encompassing normal karyotype AML patients at diagnosis before intensive chemotherapy was divided into two groups consisting of low and high scores for the VEN_plus_AraC_UP-gene signature, the high score group displayed a significantly shorter 3 years overall survival (H.R. 1.88; Fig. 4G). Thus, VEN_plus_AraC_UP-gene signature is predictive of poor prognostic in normal karyotype AML patients and associated with significantly worse patient survival. This result suggests that when therapeutic resistance is already present at diagnosis in some patients, it might represent a pleiotropic mechanism that might be predictive of the survival rate post-chemotherapy as well.

### Transcription factors MITF, E2F4, and TP53 highlighted as central drivers of resistance to duplet therapy VEN+AraC

We applied SCENIC, a robust clustering method for the identification of stable cell states from scRNA-seq data based on the underlying gene regulatory networks (GRNs)^58^. More precisely, SCENIC predicts transcription factors governing cell states and candidate transcription factor target genes, defined as regulons. Applying this method to our data, we confirmed the clear distinction of the states of VEN+AraC treated cells compared to other AML cells. Indeed, SCENIC identified two clusters VII and IX, specific for VEN+AraC treated cells (Fig. 5A) that correspond to previously Seurat identified clusters #6 and #13 respectively (Fig. 5A; Supplementary Fig. S9A). Moreover, SCENIC predicted a complex regulatory network underlying the cluster #13 (SCENIC cluster IX) involving regulons leaded by MITF, GATA1, GATA2, TAL1 (Fig. 5B-5C, Supplementary Fig. S10A-S10E). Interestingly, the three latter transcription factors are well known to be involved in transcriptional programs of hematopoiesis stem cells and hematopoiesis^59–62^, confirming the HSC-like phenotype of this cluster (Supplementary Fig. S8C). MITF is notably regulating genes involved in mitochondrial and oxidative metabolism in different cell types^63,64^, Among the common target genes that were predicted to be regulated by those TFs and composing the core of the regulatory network, numerous genes were identified as transcriptional markers (highlighted in black circle; Fig. 5C) of VEN+AraC resistance gene signature (VEN_plus_AraC_UP-gene signature, Fig. 4F). These data support the notion that MITF, GATA1, GATA2 and TAL1 might affect AML patient response through the regulation of genes involved in lipid metabolic process and inflammatory response (Supplementary Fig. 8E). Regarding cluster#6 (SCENIC cluster VII), SCENIC uncovered as top candidate BATF and p53 regulons (Fig. 5D; Supplementary Fig. S10F), both known to be involved in mitochondrial and oxidative metabolism^65,66^. Moreover, p53 and E2F4 were identified as the two common active regulons (included in top 25 of active regulons) of the three VEN+AraC-treated cell clusters (Fig. 5E; Supplementary Fig. S10G-S10J). Importantly, this study demonstrates the existence of transcriptional and phenotypic heterogeneity within the tumor derived from patient in mice BM and proposes mitochondrial metabolism and transcription factors as candidates for targeted therapy in AML.

**Figure 5.**
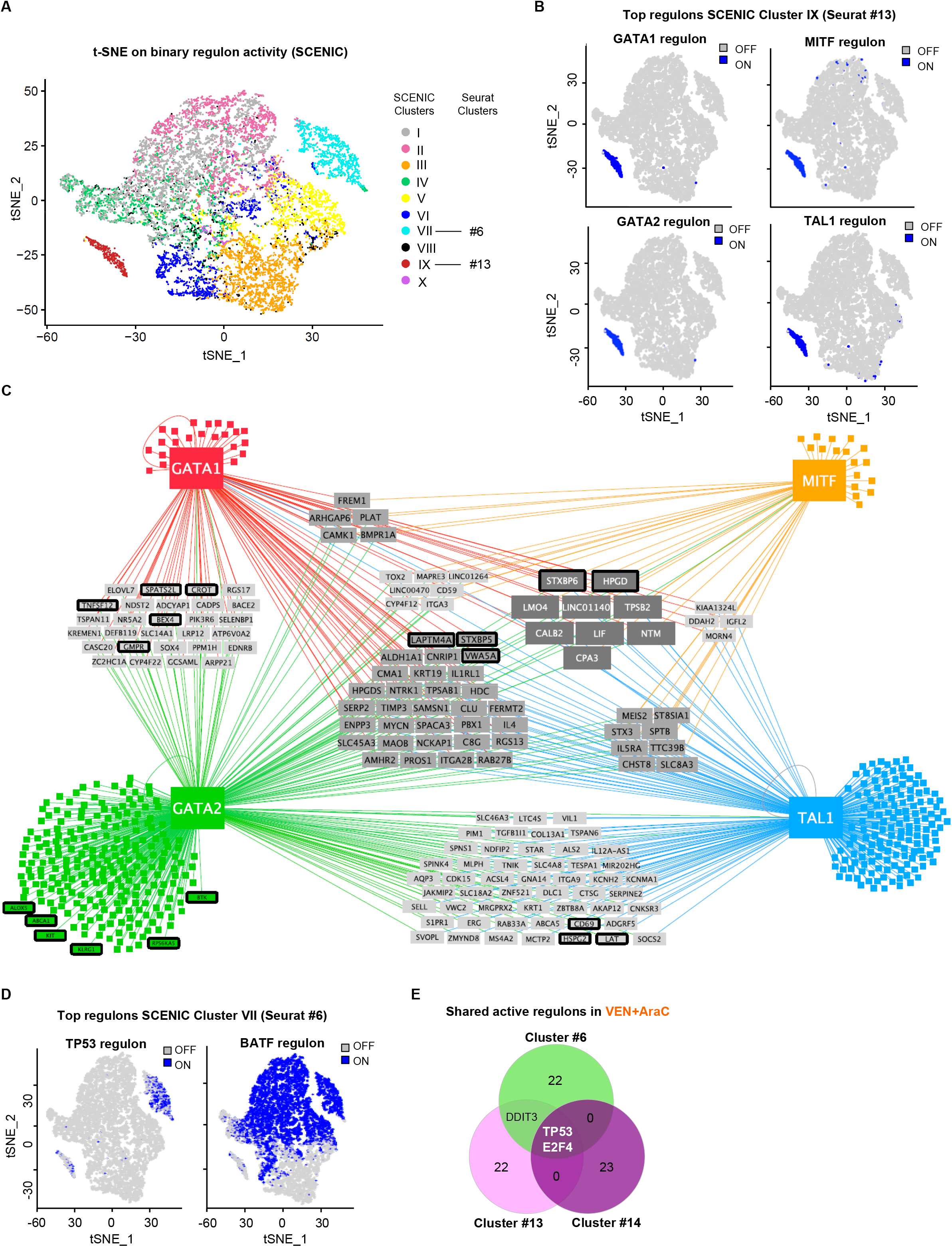
Single cell RNA-seq analysis reveals a duplet-specific regulon activity of VEN+AraC *in vivo*. **(A)** t-SNE plot displaying SCENIC clusters identified based on Gene Regulatory Network inference. Colors indicate SCENIC clusters. **(B)** t-SNE plot displaying binary activity of cluster #13 (SCENIC cluster IX) top regulons GATA1, GATA2, MITF and TAL1. **(C)** Critical nodes driving cluster #13 (SCENIC cluster IX) identified by SCENIC Gene regulatory network analysis. The top predicted TFs and their target genes are shown. Highlighted in black are the genes that were identified as clusters #13 markers in Seurat analysis. **(D)** t-SNE plot displaying binary activity of cluster #6 (SCENIC cluster VII) top regulons TP53 and BATF. **(E)** Venn Diagram displaying the shared active regulons between cluster #6, #13 and #14. TOP25 of active regulons (based on adjusted P-value) of each cluster were compared.

### Targeting ETCI in minimal residual disease from duplet therapy VEN/AraC strongly enhances time-to-relapse *in vivo*

Having shown that VEN+AraC combination is far more effective than AraC alone *in vivo* and that residual single cells exhibit transcriptional and regulatory programs associated with specific developmental, mitochondrial energetic and metabolic properties, we further identified nine most upregulated genes that are common to the three cell clusters specific for duplet residual disease *in vivo* (Fig. 6A). Interestingly, amongst these putative targets, 3 genes (NDUFA3, NDUFA13, ROMO1) encoded for mitochondrial proteins involved in NADH dehydrogenase complex assembly (Fig. 6A-6B), suggesting that ETCI activity is implicated in the duplet resistance. Gene set enrichment analysis confirmed that single cell clusters #6 and #13 (Fig. 6C) and the bulk cell population (Fig. 6D-6E) from residual disease after VEN+AraC were associated with genes for respiratory chain complex I.

**Figure 6.**
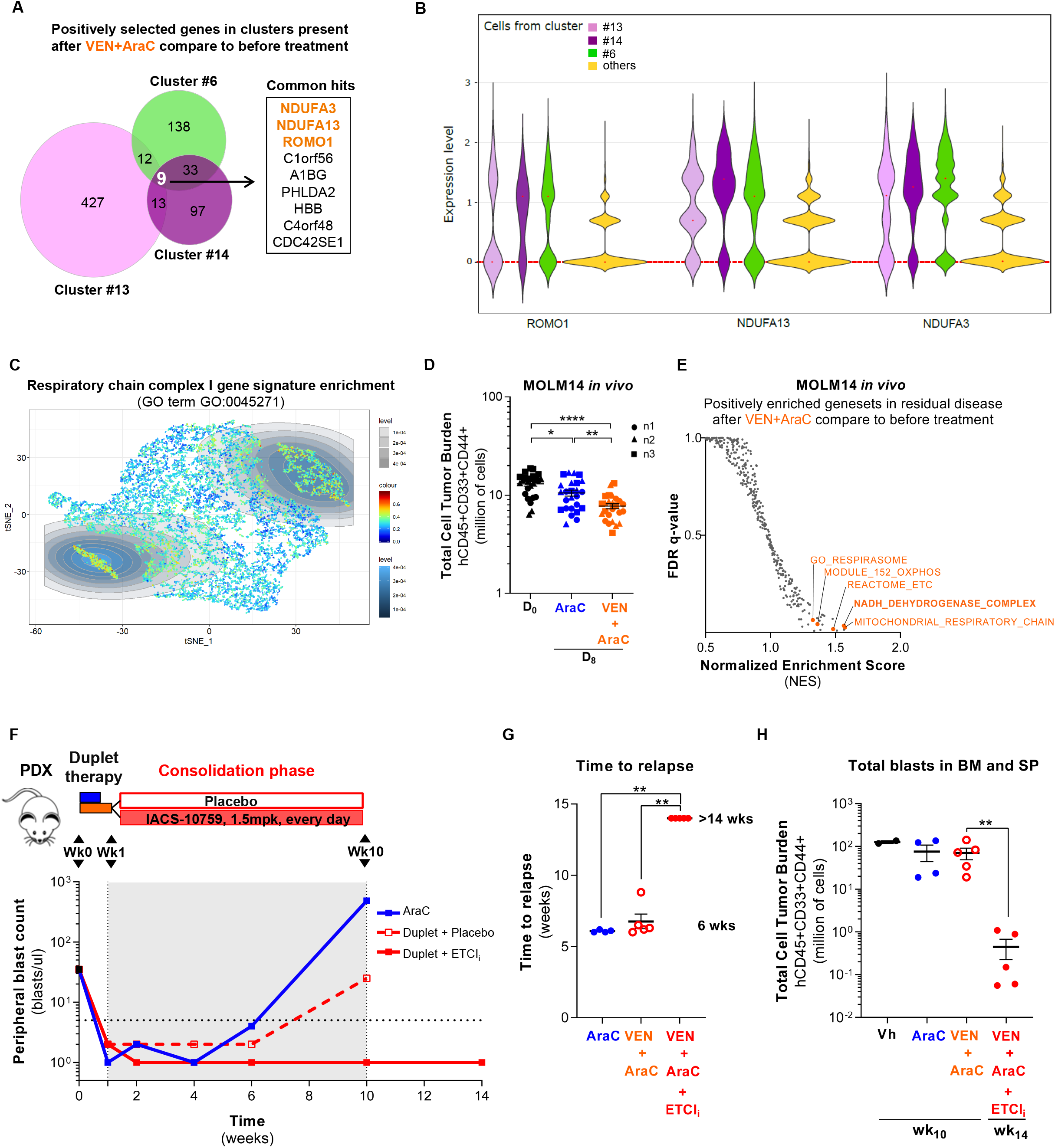
Resistance to duplet venetoclax plus cytarabine therapy is associated with ETC-I dependency in AML. **(A)** Common positively selected genes in the three clusters appearing following the duplet therapy (#6, #13, #14) *in vivo* based on single cells RNA expression levels. **(B)** Violin plots of ROMO1, NDUFA13 and NDUFA3 genes expression within clusters 6, #13, #14 and others clusters. Red dotted line indicates the mean of the dataset. **(C)** Visualization on t-SNE plot of single cell scores for the complex I gene signature using Single Cell Signature Explorer. **(D)** Cumulative total cell tumor burden of human viable CD45^+^CD33^+^CD44^+^ AML cells from Cell Line Derived Xenograft MOLM14 assessed in bone marrow and spleen at day 0 (D_0_) and at day 8 (D_8_) for the two different treatments AraC and VEN+AraC (based on the study design Fig 2A) by flow cytometry. Each dot is a mouse. Mean ± SEM of each treatment condition are represented. **(E)** Gene set enrichment analysis (GSEA) of the *in vivo* CLDX MOLM14 following the duplet therapy venetoclax plus AraC at D_8_ versus before treatment at D_0_. Positively enriched gene ontology terms. **(F)** The ETC-I inhibitor IACS-10759 was used in consolidation phase (week 1 to week 10) in PDXs after the duplet therapy (week 0 to week 1). Peripheral AML blasts in PDXs treated with the experimental arm (duplet therapy plus IACS-10759) or placebo comparator arm (duplet therapy plus placebo) were followed by peripheral blood collection until the relapse of the placebo comparator arm (duplet therapy plus placebo). The peripheral blast count of viable AML cells (CD45^+^CD33^+^CD44^+^AnnexinV-) in Patient Derived Xenograft TUH87 with the duplet therapy in association or not with the ETC-I IACS-010759 is shown. Each dot is the mean of the peripheral blast count of at least four mice. **(G)** The time to relapse of PDX TUH87 treated with AraC, VEN+AraC or VEN+AraC+IACS-10759 based on the peripheral blast count (F). Relapse was considering when the peripheral blast count overcomes the limit of detection (LoD, five blasts/μl). **(H)** The cumulative total cell tumor burden of human viable CD45^+^CD33^+^CD44^+^ AML cells assessed at the duplet therapy (placebo arm) relapse (week 10) and 4 weeks after (week 14), in mice bone marrow and spleen.

Consistent with residual disease composed of resistant AML cells crucially relying on ETC complex I, we directly tested the role of ETCI in mediating relapse *in vivo*. Thus, we propose a new rationale for a triplet therapy incorporating an ETC complex I inhibitor in consolidation phase after VEN+AraC treatment (Fig. 6F). To end this, a mouse cohort from one PDX treated with VEN+AraC was randomized in two arms, one with low dose (1mg/kg/day for 9 weeks) of IACS-010759, a new ETCI inhibitor^26^, and another with placebo. The disease was monitored by the measurement of human blasts in mice peripheral blood (PB; Fig. 6F). We showed that the time to relapse in PB was significantly higher (at least 14 weeks post-VEN+AraC) in the IACS-010759 arm compared to placebo (6 weeks post-VEN+AraC) or to AraC alone (6 weeks post- AraC; Fig. 6F-6G). Fourteen weeks post-treatment, tumor cell burden in bone marrow and spleen was two-log magnitudes of order lower in the IACS group compared to the placebo group or with AraC alone even 4 weeks before post-VEN+AraC or post-AraC at 10 weeks, respectively (Fig. 6H).

In summary, our study highlights the pivotal role of mitochondrial metabolism and ETCI dependency in therapeutic resistance and supports the design of mitochondrially/metabolically-directed therapeutic combinations in AML.

## Discussion

Growing evidence shows that energy metabolism and mitochondrial adaptation impact the development and progression of cancer. Notably, several studies have shown that mitochondrial function and OxPHOS are associated with drug resistance^8,12–14,16,18–20^. However, the mechanism by which AraC increases OxPHOS in AML cells has remained largely unknown. Our work demonstrates a major role of BCL2 in the metabolic control of cell death and apoptosis, especially in the context of AraC therapy. BCL2 affects the MOMP, mPTP opening and overall mitochondrial function in residual cells post-AraC. Previous works showed that BCL2 can be localized to both mitochondria and endoplasmic reticulum (ER) and that overexpression of BCL2 helps maintain Ca2+ homeostasis^67,68^. Importantly, mitochondrial Ca^2+^ is critical for both OxPHOS and cell death processes. Indeed, mitochondria need Ca^2+^ to provide reducing equivalents from the TCA cycle to support OXPHOS and sustain mitochondrial bioenergetics^67,69^. Elevated mitochondrial matrix Ca^2+^ stimulates four Ca2+-sensitive dehydrogenases (DH), mitochondrial FAD-α glycerol phosphate DH, pyruvate DH, isocitrate DH and α-ketoglutarate DH^70,71^. The concerted activation of these dehydrogenases by Ca2+ critically directs the activity of mitochondrial metabolism and the production of ATP^72^. Therefore, by affecting mitochondrial Ca^2+^ uptake, BCL2 also modulates OxPHOS upon AraC treatment while still protecting cells from Ca^2+^ overload and cell death, leading to closure of the mPTP.

In accordance with this mechanistic role of BCL2 in AraC resistance, treatment with VEN+AraC induces excessive mitochondrial Ca^2+^ accumulation that sensitizes mPTP opening and impairs ETC1, TCA cycle and mitochondrial respiration in a caspase-dependent manner. This induces MOMP, dissipates mitochondrial membrane potential and stimulates apoptosis, especially when matrix Ca2+ overload is accompanied by oxidative stress and adenine nucleotide depletion to enhance opening of the mPTP^72,73^. This clearly demonstrates that the mechanisms-of-action of VEN+AraC and VEN+AZA are different. While both VEN-based combinations target OxPHOS in AML, VEN+AZA suppresses it by decreasing ETCII succinate dehydrogenase glutathionylation through the inhibition of cysteine uptake and related glutathione biosynthesis. The inhibition of succinate dehydrogenase activity of ETCII selectively targets LSCs in AML patients^45^. In contrast, LSCs obtained from relapsed AML patients are not any more reliant on amino acid metabolism due to their ability to compensate through increased FA metabolism^46,53^. Interestingly, monocytic AML patients exhibited low responses to VEN+AZA caused by a reliance on MCL1 (and not BCL2) to mediate OxPHOS and survival^49,74^. Here, based on our PDX models and patient RNAseq from clinical trials comparing VEN+LDAC and VEN+HMA^25^, we discovered that the response to VEN+AraC does not correlate with developmental/differentiation status as opposed to VEN+AZA^49^. Moreover, the basal metabolic state is an important determinant of response since patients enriched in mitochondrial metabolism gene signatures are highly responsive to VEN+AraC but not to VEN+AZA. Therefore, these two VEN-based combinations are clinically mirrored and complementary. Although both VEN-based combinations increase complete remission rate in AML patients who were ineligible for intensive chemotherapy^25^, VEN+AraC was less effective than VEN+HMA in improving overall survival in this patient subgroup. However, our study demonstrates that AML patients might be better stratified for this former therapy and higher doses of AraC might be critical to improving long-term efficacy of this combination. Indeed, VEN+AraC might be used in patients relapsing or non-responsive to chemotherapy, or after failure of frontline VEN+HMA regimen (43% of patients with R/R disease, 20% with primary refractory disease; Maitis et al Haematologica 2020) when they exhibit the expression of OxPHOS markers. This knowledge will help and guide physician expectations and patient management for VEN+AraC.

Finally, resistance to both duplet therapies inevitably develops and the inclusion of a third agent will be mandatory to prolong patient survival. Several mechanisms of tolerance or resistance to VEN alone or VEN+AZA in AML have been proposed that are related to signaling pathways such as FLT3, RAS or MAPK, to bi-allelically perturbing p53^25,74^, to shifting in cellular anti-apoptotic dependencies or to mitochondrial metabolism^4^. Targeting genes involved in mitochondrial structure and function such as the heme biosynthetic pathway or the mitochondrial chaperonin *CLPB* sensitizes AML blasts to VEN or VEN+AZA combination in AML in a p53-independent manner^75,52,76–78^. To identify the mechanisms of resistance to VEN+AraC, we characterized MRD after VEN+AraC treatment *in vivo* using single-cell sequencing. Consistent with previous findings^79^, this analysis revealed that VEN+AraC MRD displayed prominent transcriptional heterogeneity and exhibited three distinct differentiation and metabolic states. However, mitochondrial NADH dehydrogenase ETC complex I was the common pathway in the three cell clusters resistant to VEN+AraC. Furthermore, using the prediction model SCENIC, we identified GATA1, GATA2, TAL1, MITF, p53, E2F4 and BATF regulons as key interrelated drivers of VEN+AraC resistance. While they are involved in AML differentiation or tumor progression, MITF, BATF, p53 regulate mitochondrial metabolism and OxPHOS^80–82^. p53 regulates several metabolic pathways including glycolysis and PPP, lipid metabolism, serine and nucleotide synthesis in cells. It also sustains the TCA cycle and fatty acid oxidation and promotes mitochondrial biogenesis by enhancing the activity of cytochrome c oxidase 2 and TFAM^80^. Strikingly, previous studies have shown that p53 controls ETC complex I and IV assembly and activity^80,83^. MITF, the other master TF found in MRD after VEN+AraC, was also shown to be involved in oxidative metabolism sustaining mitochondrial biogenesis in melanoma and maintaining the amino acid pool through the regulation of autophagy in pancreatic cancer^81,82,84^. Interestingly, E2F4, the other common active regulon of cell clusters specific to VEN+AraC, was recently shown to be involved in disease progression and was associated with a poor prognosis in AML^85^. Together, these data highlight both the crosstalk between mitochondrial metabolism and survival pathways and the metabolic control of cell death and apoptosis^78,86^, supporting the design of mitochondrial/metabolically-directed combinatory therapies in AML. Our results further confirm the emerging role of ETC complex I and its dependency in AML biology and drug response^26,87,88^. Our work also reinforces our line of reasoning that manipulating mitochondrial function with an OxPHOS inhibitor in the presence of VEN+AraC treatment *in vivo* can be effective in overcoming treatment resistance. Indeed, concomitant inhibition of BCL-2 and OxPHOS could induce synergistic cell death in AraC resistant AML cells and conceivably prevent the emergence of drug resistance to the combination of VEN+AraC.

## Methods

### Primary AML cells

Primary AML patient cells from peripheral blood have been collected during routine diagnostic procedures at the Toulouse University Hospital (TUH), after informed consent and stored at the HIMIP collection (BB-0033-00060). According to the French law, HIMIP collection has been declared to the Ministry of Higher Education and Research (DC 2008-307 collection 1) and obtained a transfer agreement (AC 2008-129) after approbation by the “Comité de Protection des Personnes Sud-Ouest et Outremer II” (ethical committee). Clinical and biological annotations of the samples have been declared to the CNIL (“Comité National Informatique et Libertés”; i.e. “Data processing and Liberties National Committee”). Peripheral blood and bone marrow samples were frozen in fetal calf serum with 10% DMSO and stored in liquid nitrogen. Human primary AML cell were grown at 37°C with 5% CO2 in Iscove’s Modified Dulbecco’s Media with Glutamax (IMDM, Gibco, Life Technologies) supplemented with 20% fetal calf serum (Invitrogen, Carlsbad, CA, USA) and 100□units per ml of penicillin and 100□μg/ml of streptomycin.

### AML Cell lines

U937 was purchased from ATCC (Manassas, VA, USA) in January 2014. MOLM-14 was obtained from Pr. Martin Carroll (University of Pennsylvania, Philadelphia, USA) and MOLM-14 and OCI-AML3 were purchased from the Leibniz Institute DSMZ-German Collection of Microorganisms and Cell Cultures (Leibniz, Germany). DSMZ and ATCC cell banks provides authenticated cell lines by cytochrome C oxidase I gene (COI) analysis and short tandem repeat (STR) profiling. Clinical and mutational features of our AML cell lines are described in Supplementary Table S1. These cell lines have been routinely tested for Mycoplasma contamination in the laboratory. Human AML cell lines were grown at 37°C with 5% CO2 in minimum essential medium-α medium with Glutamax (MEMα, Gibco, Life Technologies) supplemented with 10% fetal calf serum (Invitrogen, Carlsbad, CA, USA) and 100□units per ml of penicillin and 100□μg/ml of streptomycin. The cultured cells were split every 2-3 days and maintained in an exponential growth phase.

### AML mouse xenograft model

Animals were used for transplantation of AML cell lines or primary AML cells in accordance with a protocol reviewed and approved by the Institutional Animal Care and Use Committee of Région Midi-Pyrénées (France). NOD/LtSz-SCID/IL-2Rγchain null (NSG) mice were produced at the Genotoul Anexplo platform at Toulouse (France) using breeders obtained from Charles River Laboratories. Mice were housed in sterile conditions using HEPA-filtered microisolators and fed with irradiated food and sterile water. MOLM-14 cells were injected into mice for a Cell Line Derived Xenograft (CLDX) and primary AML cells were injected for a Patient Derived Xenograft (PDX) as previously reported^43^. Briefly, mice (6-9 weeks old) were sublethally treated with 20 mg/kg of busulfan for CLDXs and 30 mg/kg for PDXs. 24 hours after, AML cells were washed in PBS, and suspended in Hank’s Balanced Salt Solution (HBSS) at a final concentration of 2×10^6^ cells per 200μL of HBSS per mouse for tail vein injection. All experiments include male and female mice and experimental groups were randomly assigned. CLDXs transplanted with MOLM-14 cell line were treated at day 10 post-transplantation for 1 week and sacrificed at day 17. The engraftment of primary AML cells in mice bone marrow and spleen is followed by peripheral blood analysis or bone marrow aspirates and took 12 to 20 weeks. Once PDXs were engrafted they were treated one week and sacrificed to determine the total cell tumor burden (TCTB) in mice bone marrow and spleen. The treatment consists in 5 days of AraC treatment (10 mg/kg for CLDXs and 30 mg/kg for PDXs) by intra-peritoneal injection, in association with 7 days of venetoclax treatment (100 mg/kg) by oral gavage. Venetoclax was solubilized in corn oil with 30% PEG400 and 10% ethanol before administration to mice. For the tri-therapy experiment, PDXs were treated with AraC plus venetoclax for one week as previously, following with a 9 weeks’ consolidation phase with a low dose IACS-10759 treatment (1.5 mg/kg) or a placebo treatment every day by oral gavage. Mice relapses were followed periodically by peripheral blood analysis. Mice were sacrificed when the placebo arm relapsed and the TCTB were assessed. IACS-10759 was solubilized in water containing 0.5% methylcellulose 400 cp before administration to mice.

### Statistical analyses

We assessed the statistical analysis of the difference between two sets of data using non-parametric Mann-Whitney test one-way or two-way (GraphPad Prism, GraphPad). P-values of less than 0.05 were considered to be significant (* P<0.05, ** P<0.01 and *** P<0.001). Detailed information of each test is in the figure legends.

## Acknowledgements

We thank all members of mice core facilities (UMS006, ANEXPLO, Inserm, Toulouse) in particular Marie Lulka, Christine Campi, Pauline Challies, Pauline Debas, Massimiliano Bardotti and Sarah Gandarillas for their support and technical assistance, and Prof. Véronique De Mas and Eric Delabesse for the management of the Biobank BRC-HIMIP (Biological Resources Centres-Inserm Midi-Pyrénées “Cytothèque des hémopathies malignes”) that is supported by CAPTOR (Cancer Pharmacology of Toulouse-Oncopole and Région). We thank Astrid Melotti and IGE3 genomic platform (Geneve University) as well as Marie Tolosini, Frédéric Pont, Frédéric Lopez (Pôle Technologique du CRCT, Inserm/U1037, Toulouse) and Frédéric Martins (GET, GENOToul, Toulouse) for bulk DNA/RNA sequencing and single cell RNA seq procedures, respectively. We are grateful to the genotoul bioinformatics platform Toulouse Midi-Pyrenees (Bioinfo Genotoul) for providing computing resources. This work was granted access to the HPC resources of CALMIP supercomputing center under the allocation 2019-T19001. Metabolomic experiments were carried out at the MetaToul-MetaboHUB core facility (National Infrastructure of Metabolomics and Fluxomics, Toulouse, France) under the supervision of Drs. Lindsay Peyriga, Floriant Bellvert and Jean-Charles Portais. MetaToul (Metabolomics & Fluxomics Facitilies, Toulouse, France, www.metatoul.fr) and LEMM are part of the national infrastructure MetaboHUB-ANR-11-INBS-0010 (The French National infrastructure for metabolomics and fluxomics, www.metabohub.fr). MetaToul is supported by grants from the Région Midi-Pyrénées, the European Regional Development Fund, the SICOVAL, the Infrastructures en Biologie Santé et Agronomie (IBiSa, France), the Centre National de la Recherche Scientifique (CNRS) and the Institut National de la Recherche Agronomique (INRA). We thank Anne-Marie Benot, Muriel Serthelon and Stéphanie Nevouet for their daily help about the administrative and financial management of our Team. The authors also thank Dr. Nathalie Mazure for fruitful discussion about mitochondrial VDAC, Prof. Marina Konopleva, Ian Majewski and Andrew Wei for the access to RNA sequencing from AML patients, and Dr Mary Selak for critical reading of the manuscript.

